# Comparative study of pre- and post-mortem perfusion of fixative for the quality of neuronal tissue preparation

**DOI:** 10.1101/2024.09.19.613859

**Authors:** Géraldine Meyer-Dilhet, Salma Ellouze, Olivier Raineteau, Julien Courchet

**Affiliations:** Univ Lyon, Univ Lyon 1, CNRS, INSERM, Physiopathologie et Génétique du Neurone et du Muscle, UMR5261, U1315, Institut NeuroMyoGène, 69008 Lyon, France; Univ Lyon, Université Claude Bernard Lyon 1, Inserm, Stem Cell and Brain Research Institute U1208, 69500 Bron, France

## Abstract

Transcardiac perfusion of fixative agent is generally recommended for quality preparations for cerebral histology, ensuring rapid and deep penetration in the tissue to preserve the most fragile brain structures. Despite being performed under anesthesia and with proper analgesia, this procedure is cumbersome for the experimenter and raises ethical questions. Recently, alternative protocols have been proposed, based on prior sacrifice of the animal followed by an injection of a fixative agent into the circulation. These so-called post-mortem perfusion protocols should in theory ensure an equivalent quality of tissue fixation, without exposing live animals to a procedure. Before adopting this new method, it is necessary to validate that sample quality is equivalent, ensuring the validity of scientific results. We performed a parallel comparison of several protocols of tissue fixation, by transcardiac or post-mortem perfusion, and measured the impact on the maintenance of axonal structures, dendritic spines, and mitochondrial morphology. Our results showed that histological parameters show variable sensitivity to perfusion condition and fixative used. For instance, axon fragmentation and altered mitochondrial morphology were observed in post-mortem perfusion groups. We furthermore determined that fixation condition had a variable effect on immunostaining, impacting detected expression level or pattern. Our results serve as a guide to orient the experimenter in selecting the best condition for optimal tissue fixation, which minimizes animal suffering while guaranteeing the integrity of the biological results obtained.

## INTRODUCTION

The use of live animals in life sciences is a major societal challenge and source of debate within the scientific community. Research based on animal has led to major advances in knowledge in fundamental and biomedical sciences and is currently essential, despite the rapid development of alternative methods. Humane research on animal adheres to the ‘3Rs’ principles, standing for ‘Reduce, Replace, Refine’, first defined in 1959. This principle is embedded in the European legislation on the protection of animals used for scientific purposes (Directive 2010/63/EU). High standards of animal care and experimentation not only benefit animals, but they are also widely beneficial for workers in contact with animal who may suffer from compassion fatigue.

Refinement is defined as a way to minimize pain, distress and harm from procedures on animals, when these procedures cannot be avoided. This is often the case in neurosciences, which concerned nearly 849 000 animals in the European Union in 2020, mostly mice and rats. These species are widely used because the complexity, organization and regulation of neural networks matches the human nervous system, allowing to elucidate fundamental brain processes that may be impaired in neurodevelopmental, neuropsychiatric and neurodegenerative conditions. Because much of the function of the nervous system relies on the study of intercellular connectivity, histological preparations are a common practice. Tissue fixation is a critical step to prevent the autolysis and degradation of tissue and tissue components that rapidly happens following the death of the animal^1^. Immersion in a solution containing fixative agent is an efficient method for fixation of small elements such as the sciatic nerve^2^. However, this method is generally considered as not appropriate for the fixation of larger tissues such as the whole brain of a rodent, because the penetration of fixative agents in the depth of tissue is too slow to prevent hypoxia and cellular changes in deeper brain structures. Transcardiac perfusion of a fixative agent is generally performed to clear blood and preserve deep cellular structures^3,4^. Transcardiac perfusion is usually performed in deeply anesthetized animals, as a terminal procedure during which the last heart beats help propel fixative into the general circulation. However, recent reports demonstrated that the choice of anesthetic and fixative agent can have a major impact on some cellular structures, such as mitochondria, with consequence on data analysis^5^. Furthermore, transcardiac perfusion has been recently challenged, and it is now proposed that perfusion could be performed post-mortem, ensuring similar tissue quality with less impact on the animal. In this study, we compared the impact of different protocols for tissue fixation by transcardiac perfusion or post-mortem perfusion on several parameters that are routinely analyzed in neuroscience studies in the mouse brain. Following in utero cortical electroporations (IUCE) of plasmids^6^, pups were randomized into four groups to compare two distinct protocols of transcardiac perfusion in anesthetized animals (with volatile and injectable anesthetic, respectively), one protocol of post-mortem perfusion, and one condition of direct immersion in the fixative agent without perfusion. Tissues were prepared in parallel, and we quantified tissue quality, axon structure, dendritic spines density and morphology, and dendritic mitochondria integrity. Furthermore, we tested distinct immunostainings. Overall, the distinct parameters in the study were unequally affected by fixation conditions. While some parameters were essentially similar regardless of the fixation condition (e.g. axon branching density and dendritic spines density), other parameters were strongly variable from one group to the other (e.g. fragmentation of axons, dendritic mitochondria integrity and phospho-CREB immunoreactivity). Overall, our results demonstrate that the choice of fixation condition can have a major impact on the quality of results and must be balanced with the impact to animal to ensure robust results.

## RESULTS

### Study design and fixation of tissues

To compare the influence of fixation condition on various parameters relevant for neurosciences, we performed *in utero* cortical electroporations (IUCE) with plasmids encoding the red fluorescent protein mScarlet-I, as well as the mitochondrial marker OMM-GFP (**Fig. 1A**). At birth, we selected pups based on transcranial fluorescence observation, in order to keep only electroporated pups for the rest of the procedure. At postnatal day (P)21, the pups were divided into four groups corresponding to the experimental conditions. Groups were equilibrated in males and females, and pups coming from different litters were split into distinct groups, to avoid a potential batch effect. All mice received an injection of buprenorphine between 30 minutes to 2 hours before starting the procedure.

**Figure 1:**
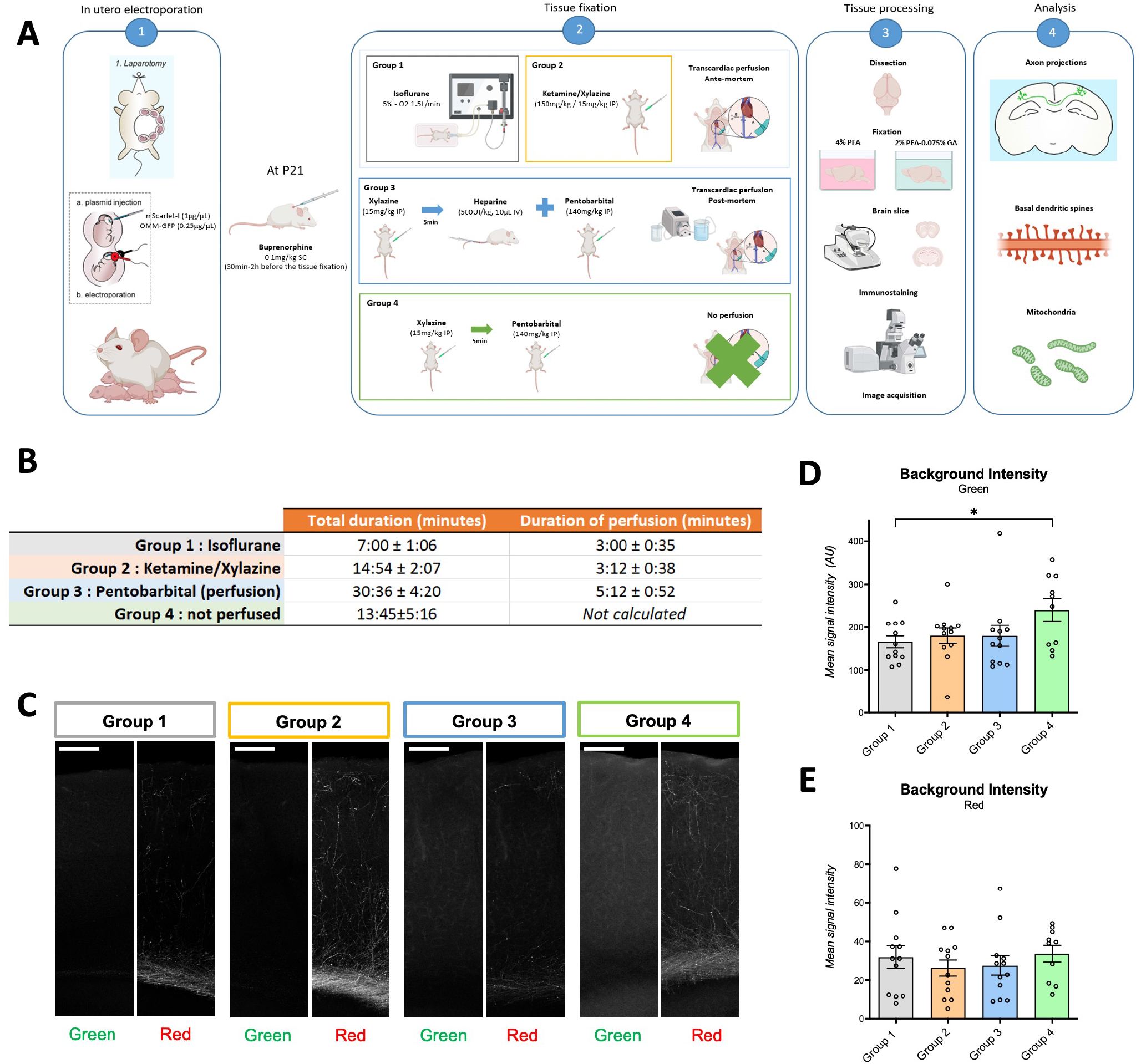
Experimental design and overall quality of tissues. **(A)** Experimental design. **(B)** Duration of the procedure and perfusion time for each group. Data are the average of 9 animals per group ± IC95%. **(C)** Detail (representative images) of confocal images acquired from coronal sections of brains for each group. Images in the green (anti-GFP) and red (mScarlet-I) channels. All images are taken with the same confocal settings (laser power and gain). Images were taken from the contralateral side (opposite to the electroporation side). Scale bar: 200μm. **(D-E)** Background signal intensity was quantified in the cortex by drawing ROIs and measuring average signal intensity (arbitrary units). ROIs were drawn in regions devoid of specific signal (electroporated neurons). Each circle on the graph represents the average of 5 ROIs taken from one image, one to four images were quantified per animal. N=10 (group 4) to 12 (groups 1, 2 and 3). Statistical analysis: one-way ANOVA with Dunnett”s multiple comparison.

Two groups (group 1 and group 2, **Fig. 1A**) corresponded to an ante-mortem transcardiac perfusion of fixative in anesthetized mice (either with volatile anesthesia for group 1, or with fixed anesthesia for group 2). Once a deep sedation state was constated (no reflexes to pinching, slow respiration), a thoracotomy exposed the heart in which the fixative agent was injected in the left ventricle. The last heart beats immediately followed fixative injection. In comparison, group 3 corresponded to a post-mortem procedure in mice that were previously euthanized (**Fig. 1A**). We first performed a light sedation, followed by intravenous (IV) injection of heparine and injection of an euthanizing agent. Transcardiac perfusion was performed post-mortem, which we defined as a respiratory and cardiac arrest. Here, we used a peristaltic pump to force the fixative agent into the general circulation, in spite of the lack of heat beats. In group 4, after an intraperitoneal (IP) injection of xylazine, mice were euthanized by pentobarbital and the brain was dipped into a fixative solution, without prior perfusion (**Fig. 1A**). This condition was designed as a reference to objectivate the need to inject fixative agent in the general circulation for good tissue quality. Hence, it was expected that groups 1-3 should perform as well or better than group 4 for all parameters analyzed.

### Implementation of the protocol and general tissue quality

Volatile anesthesia was the fastest way to reach deep sedation (**Fig. 1B**). Methods based on injections took longer time, during which mice were put in a quiet box for sedation to take place. When considering only the perfusion, the procedure was overall faster in groups 1 and 2, where heart beat helped the delivery of the fixative agent. Although all groups were subject to at least one injection (buprenorphine prior to the procedure for proper analgesia), groups required different level of handling of the animals, since each injection required some contention. The most handling and contention was required for animals from group 3, who in total received 4 injections.

For perfused animals (groups 1-3), rigor of the limbs and tail was indicative of a good fixation. The discoloration of the brain caused by blood removal was a further sign that perfusion was efficient (**Suppl. Fig. 1**), although we noticed some variability in the quantity of residual blood in group 3. In comparison, blood was still present in the brain of non-perfused animals.

Following the procedure, brains were dissected and post-fixed with 4% Paraformaldehyde (PFA) overnight (**Fig. 1A**). We then performed coronal sections, immunostainings and confocal imaging. We first reasoned that perfusion quality might change background fluorescence of the tissue. Overall, background fluorescence was more important in the green channel, which is typically more prone to autofluorescence. Measurement of average fluorescence level in the cortex revealed as expected that non-perfused brains (group 4) had more background fluorescence than other conditions (**Fig. 1C-D**). Groups 1, 2 and 3 had similar levels of background fluorescence, although there was some variability between animals revealing potential heterogeneity in perfusion quality. On average, background in the red channel was much lower than in the green channel, thus individual differences in background signal had a lesser overall impact (**Fig. 1E**).

### Sub-optimal perfusion conditions are associated with axon fragmentation and can affect the quantification of axonal projections

We first sought to assess the impact of fixation conditions on the more fragile neuronal structures. Axons are long, thin structures prone to rapid degradation upon cellular stress. Neurons from the superficial layer of the somatosensory cortex typically project axons to cortical layers II/III and V of the ipsilateral and contralateral cortex (**Fig. 2A**). Because all animals came from the same electroporation procedure and were allotted randomly, our prediction was that any difference measured on average on axon branching patterns should be the result of the fixation condition. On ipsilateral layer V, the dense ramification of axons can be assessed by measuring the optical density in the red channel, corresponding to the fluorescence of mScarlet-I by electroporated neurons. Indeed, there was no statistically significant differences between groups in the axon density measured in the ipsilateral cortex after background fluorescence subtraction (one way ANOVA test P=0.385 for an effect of group), indicating that the signal to noise ratio was correct in the ipsilateral cortex (**Fig. 2B**). In contrast, we noted differences between groups in the contralateral hemisphere. The best results were observed with group 1, where there was a marked density of axon projections in the S1/S2 boundary in the somatosensory cortex, with axons that were well marked (**Fig. 2C**, left). In comparison, terminal axons density was reduced in groups 2 and 3, and strongly reduced in group 4. Interestingly, we noted that axons appeared more fragmented in groups 3 and 4, which may in part explain the apparent decrease of axon projection density (**Fig. 2D**). A two-way ANOVA test confirmed an overall effect of genotype (F(3, 44)=5.190, P=0.0037) on axon density profiles. Multiple comparison assays revealed a decrease in the density of axons on cortical layers II/III and V for groups 2 and 3 as compared to group 1, despite a similar density of afferent axons in the white matter (WM) (**Fig. 2E**). The worse condition being in the non-perfused group (group 4).

**Figure 2:**
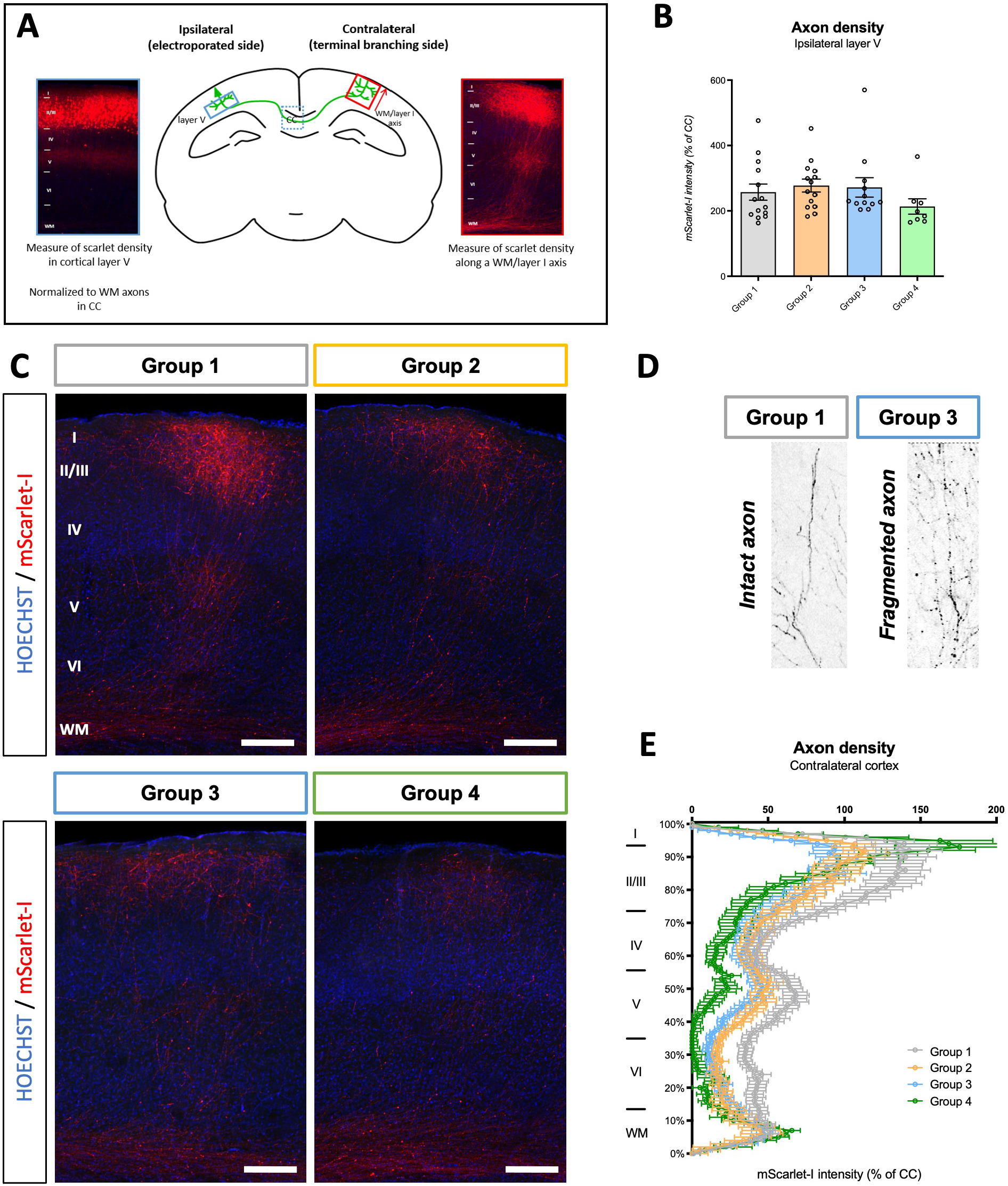
Effect of perfusion condition on axons. **(A)** Representation of a coronal brain section showing the soma of electroporated neurons (left side, ‘ipsilateral’) and axon projection and pattern of branching on the ipsilateral and contralateral cortex. The method of quantification of axon branching varies between the ipsilateral and contralateral side to adapt to the density of projections. Normalization to the density of white matter axons (WM) in the corpus callosum (CC) serves as a way to compensate for potential differences in the number of electroporated neurons. **(B)** Quantification of axon density in layer V of the ipsilateral cortex. Data represents the average signal value in 5 ROIs (arbitrary units), and normalized to signal intensity measured in the CC. Each circle on the graph represents the average value for one brain slice, two slices were analyzed per animal. N=8 (group 4) to 14 (groups 1, 2 and 3). Statistical analysis: one-way ANOVA (F(3,44)=1.039, p=0.385). **(C)** Representative images of terminal axon branching on the contralateral side. Scale bar: 200μm. WM: white matter. I, II, III, IV, V and VI: cortical layer identity. **(D)** Magnification (x4) shows axon fragmentation in the case of group 3, not group 1. **(E)** Quantification of mScarlet-I intensity along a white matter to layer I axis in the distinct groups. Data represents the average value for N=6 (group 4), 13 (group 3) to 16 (groups 1, 2) brain slices from 4 to 6 animals. Statistical analysis: two-way ANOVA.

### Fixation conditions had a moderate impact on dendritic spines density and morphology

Synapses are the focus of numerous studies owing to their central importance in brain function. In cortical pyramidal neurons, glutamatergic synapses are formed at the extremity of dendritic spines whose density and shape dynamically changes, reflecting neuronal activity. We reasoned that sub-optimal tissue fixation could alter the detection of dendritic spines, making it harder to count or classify them. Moreover, dendritic spines are dynamic structures which could be rapidly altered post-mortem if fixation is not optimal. Thus, we imaged isolated basal dendritic segments from electroporated cortical neurons and manually analyzed dendritic spines (**Fig. 3A-B**). We first reasoned that fixation condition may affect our capacity to detect dendritic spines, or that slow fixation of synaptic structures could lead to dendritic spines shrinkage. Thus, we first compared dendritic spines density, measured from at least 4 basal dendritic segments from 4 individual neurons for each animal (**Fig. 3C**). After quantification, there was no overall effect of the group on the density of dendritic spines (F(4,72)=0.8222, p=0.4861) (**Fig. 3E**). It has been proposed that glutaraldehyde in combination with paraformaldehyde contributes to better prepare tissues for synapses and mitochondria^5^. Thus, we compared results using an alternative 2% paraformaldehyde-0.075% glutaraldehyde solution (PFA-GA) instead of 4% paraformaldehyde (PFA) (**Fig. 3D**). Yet as for PFA, the density of dendritic spines was similar in all 4 groups upon fixation with PFA-GA (F(4,111)=0.5654, p=0.639) (**Fig. 3F**). Pairwise comparisons confirmed that no dataset were statistically different from one another. Interestingly, we noted a trend toward a global decrease of dendritic spines density in all conditions after fixation with PFA-GA, which likely comes from the overall increased autofluorescence of the tissue caused by glutaraldehyde ^7^.

**Figure 3:**
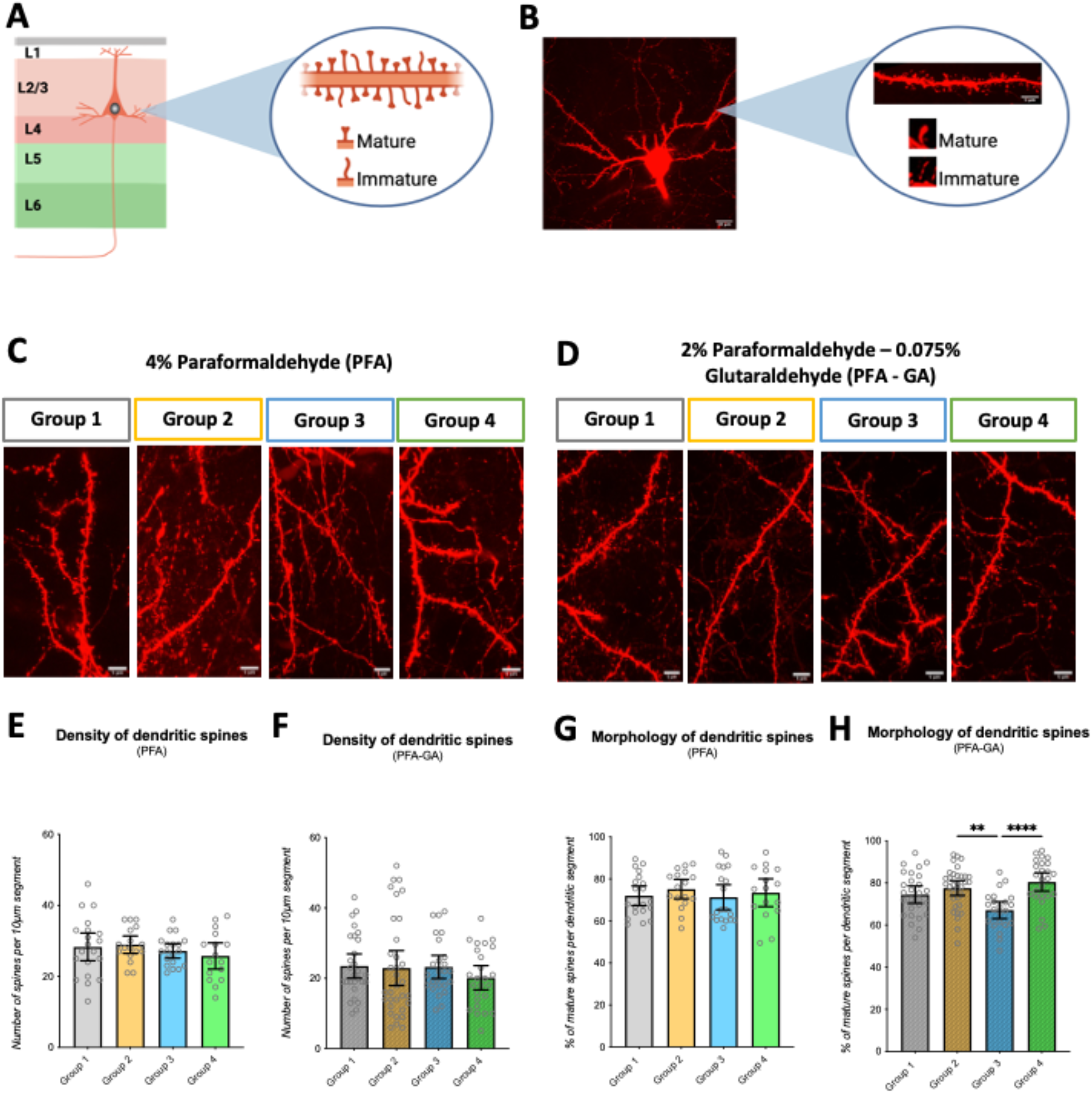
Effect of perfusion condition on basal dendritic spines. **(A)** Schematic representation of the experimental design, and **(B)** example of an optically-isolated neuron, from which segments of basal dendrites could be isolated and analyzed. Spines were classified into immature or mature based on their morphology. **(C)** Typical examples of dendritic segments from the different groups upon fixation with PFA. Scale bar 5μm. **(D)** Typical examples of dendritic segments from the different groups upon fixation with PFA-GA. Scale bar 5μm. **(E-F)** Quantification of dendritic spines density in each group upon fixation with PFA or PFA-GA. **(G-H)** Quantification of mature dendritic spines expressed as a percentage of total spines upon fixation with PFA or PFA-GA. All quantifications were performed blind to condition. N (animals) = 5 for groups 1, 2 and 3, and N (animals) = 4 for group 4. For each animal, we imaged and analyzed dendritic segments of 4 neurons. Each dot represents a neuron.

In parallel, we reasoned that the speed of tissued fixation might have a bigger impact on spine morphology, since plasticity happens on a faster time scale than spine addition/elimination^8,9^. Dendritic spines were sorted into matures (larger head, ‘mushroom’ shaped) and immatures (filopodial or long necked) and compared groups (**Fig. 3A-B**). Although as expected from animals at this age, we observed on average a majority of mature spines in all conditions, a statistical analysis demonstrated a group-effect for both the PFA (F(4,106)=12.03, p<0.0001) and PFA-GA condition (F(4,107)=7.785, p<0.0001) (**Fig. 3G-H**). Pairwise comparison revealed a trend toward more mature spines in group 2, which could be due to the dissociative effect of ketamine and its compound effect on NMDA receptors in excitatory and inhibitory cortical neurons^10,11^. Conversely, mature spines were reduced in group 3, which could be caused by the overall longer anesthesia time before tissue fixation. Overall, our data reveal that tissue fixation condition may have a small, yet significant effect on dendritic spines.

### Post-mortem fixation is associated to altered morphology of mitochondria

Mitochondria undergo rapid cycles of fusion and fission in response to cellular environment. Mitochondrial length is thought to be correlated with their metabolic activity, and furthermore mitochondria morphology is associated to pathological states^12^. Hence, fixation conditions should preserve mitochondria morphology for accurate research on neuronal energy metabolism. Previous studies demonstrated that peri-mortem hypoxia has a major impact on the quality of mitochondria preparation^5^. Here, we measured the impact of fixation on dendritic mitochondria morphology. As a whole, mitochondria appeared more elongated in groups 1 and 2, whereas mitochondria were overall shorter in group 3, and especially in group 4 (**Fig. 4A-B**). Furthermore, we observed frequent swollen mitochondria, suggesting they were actively being fragmented at the time of fixation, in neurons from group 3 and 4 (**Fig. 4C**). Quantification of average mitochondria length per dendritic segment confirmed an overall effect of the experimental condition on mitochondria length in PFA-fixed animals (one-way ANOVA F(3,103)=3.561, p=0.017) (**Fig. 4D**). Pairwise comparison revealed a decrease of the average mitochondria length in group 3 and 4 neurons when compared to group 1 or group 2. A similar result was obtained when considering the cumulative length of mitochondria regardless of the neuron or animal or origin (**Fig. 4E**). Interestingly, the overall same group effect was observed in the PFA-GA fixed neurons (**Fig. 4F-G**). Compared to the PFA-fixed conditions, there was less data variability in the PFA-GA samples, although mitochondria were overall smaller in size, unlike recent reports that glutaraldehyde fixation protects mitochondria from fragmentation^5^. Collectively, our data demonstrates that perfusion and tissue fixation have a major effect on the morphology of dendritic mitochondria.

**Figure 4:**
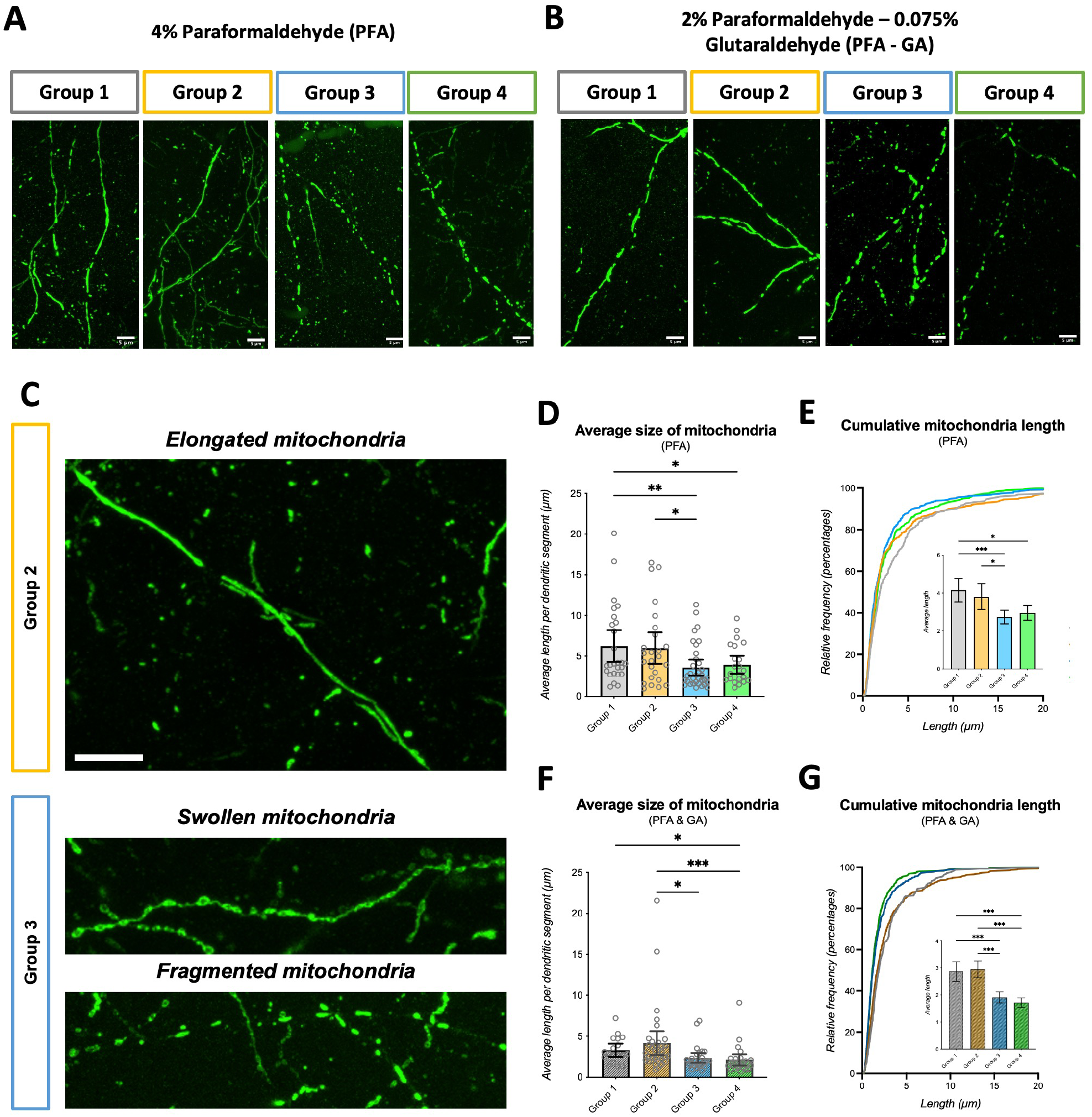
Effect of perfusion condition on dendritic mitochondria. **(A)** Detail (representative images) of dendritic mitochondria observed by confocal microscopy in each group, following perfusion/fixation with PFA, **(B)** and with PFA-GA. Scale bar 5μm. **(C)** Examples of elongated, swollen and fragmented mitochondria observed in dendritic segments. Scale bar 5μm. **(D)** Measurement of the average length of mitochondria per dendritic segment upon fixation with PFA. Data represents the average length of mitochondria in one dendritic segment. Each circle on the graph represents one dendritic segment. N=22 to 34 segments per group. Statistical analysis: one-way ANOVA (F(3,103)=3.561, p=0.017). **(E)** Cumulative frequency distribution of mitochondria size per group, and average mitochondria length upon fixation with PFA. Insert shows the average length of all mitochondria. N=354 to 525 mitochondria. Statistical analysis: one-way ANOVA (F(3,1665)=6.984, p<0.0001). **(F)** Measurement of the average length of mitochondria per dendritic segment upon fixation with PFA-GA. Data represents the average length of mitochondria in one dendritic segment. Each circle on the graph represents one dendritic segment. N=18 to 33 segments per group. Statistical analysis: one-way ANOVA (F(3,97)=3.552, p=0.017). **(G)** Cumulative frequency distribution of mitochondria size per group, and average mitochondria length upon fixation with PFA-GA. Insert shows the average length of all mitochondria. N=240 to 562 mitochondria. Statistical analysis: one-way ANOVA (F(3,1828)=24.8, p<0.001).

### Immunochemistry is known to depend upon fixation quality

Finally, we wanted to see the impact of the fixation condition on histological preparations. For this we used various classic antibodies which we know by experience might be more or less sensitive to tissue fixation conditions. We first tested if distinct fixation condition could impact cell integrity, which could alter the quality of tissues and lead to artifacts. We thus evaluated neuronal apoptosis, an intrinsic program leading to neuronal cell death, which occurs during brain development and in pathological context (neurodegenerative diseases)^13^. To do that, we performed cleaved caspase-3 immunostaining (**Fig 5A** and **Suppl. Fig. 3A**). We saw a few apoptotic neurons present on the histological sections, especially in the hippocampus, which demonstrates that the cleaved caspase-3 staining worked properly. However, there was no notable difference regardless of the experimental group observed, ruling out that the initiation of a stress response leads to apoptotic cell death, probably because the time of fixation remains fairly limited after all.

**Figure 5:**
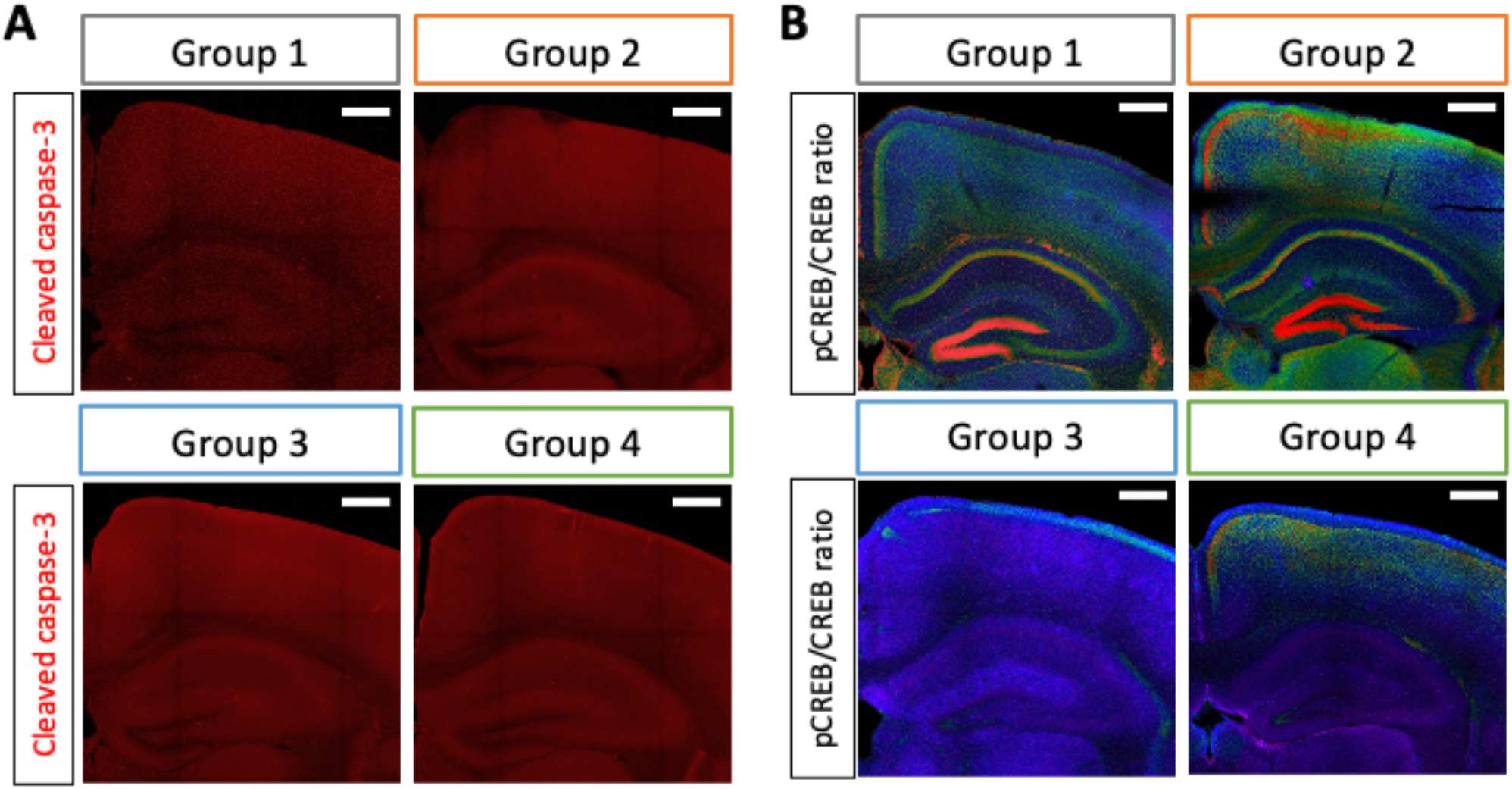
Effect of perfusion condition on axons on immunostaining. **(A)** Neuronal apoptosis observed by cleaved caspase-3 staining in each group, following perfusion/fixation with PFA. **(B)** Ratio between phosphorylated CREB (pCREB, Ser133) and non phosphorylated CREB in each group. Images are colorized on a rainbow LUT (purple: low ratio. red: high ratio). Magnification of the cortex and hippocampus. Raw images for CREB and pCREB provided in **Suppl. Fig. 4**. Scale bar: 500μm.

We next examined CREB (Ca/cAMP responsive element binding protein), a transcription factor involved in processes underlying neuronal plasticity leading to learning and memory. CREB is a so-called immediate early gene, whose rapid phosphorylation on serine 133 and activation occurs upon elevation of cAMP or CA^2+^ in the cytoplasm and this is usually correlated with the long-term memory or after a psychophysical stress^14–16^. For this staining, we observed marked differences between groups. Groups 1 and 2 showed some pCREB immunoreactivity which corresponded to expected patterns in the mouse brain, with nuclear signal in several brain regions, strong in the cerebral cortex and the hippocampus, and with the strongest signal in the dentate gyrus (**Fig. 5B** and **Suppl. Fig. 3B**). pCREB signal was overall stronger in group 2 than in group 1, and the pCREB/CREB ratio was slightly different in some brain regions between the two groups (see for example the cingulate cortex or the CA3 region of the hippocampus), which may correspond to the effect of distinct anesthetic agents. In contrary, the pCREB signal was strongly reduced in group 3 which presented the lowest signal overall. In group 4, some pCREB signal comparable to group 2 was observed in the superficial part of the cortex but rapidly signal extinguished in deeper brain regions, which most likely corresponds to the slow penetration of fixative deep into tissues. Observations deeper in the tissue in areas of the brain known to be involved in memory such as hippocampus (especially dentate gyrus) demonstrated a profound loss of pCREB staining which was weak or even non-existent in groups 3 and 4 compared to groups 1 and 2 (**Fig. 5B**). Although we cannot determine from this data if fixation conditions lead to artefactual extinction of CREB activity or if the immunostainings are affected by sample preparation or both, we conclude that the fixation conditions from groups 3 and 4 were not suitable for the study of CREB staining.

## DISCUSSION

In this study, we compared three methods for brain samples preparation. Methods 1 and 2 rely on the ante-mortem injection of a fixative agent in the general circulation by transcardiac perfusion in an anesthetized animal. Method 3 relies on the post-mortem delivery of a fixative agent, and relies on a peristaltic pump to distribute the fixative agent in the circulation of an animal that has been put to sleep. A 4^th^ group, where brains were fixed by simple immersion in the fixative solution, served as reference to measure objective benefits of methods 1-3. Brains were subsequently processed and we analyzed parameters that would be typical for such experiment, such as axon projections, dendritic spines density and morphology, mitochondria morphology, or immunostaining. Importantly, our study demonstrates that the method of tissue fixation has a major impact on experimental parameters, warranting caution when comparing results that were not obtained with the same method. Furthermore, there was not clearly one condition that gave the best results for each and every analysis parameter. Rather, our results suggest that tissue fixation and sample preparation must be adapted to the needs of the experiment that will be performed afterwards. As a general rule, the more dynamic a biological process is, the more sensitive it is to fixation protocol. This highlights the importance or pre-defined analysis parameter, as well as of pilot studies, to ensure that the most valid scientific information may be obtained from each experiment. It is assumed that brain structures deteriorate within minutes following the animal”s death. Accordingly, our results demonstrate that the method of fixation influences the axonal structure and is critical for preserving mitochondrial morphology. Although the causes of these changes are multiple, one possible explanation could be the spreading depolarization waves and excitotoxicity due to neurotransmitter release^17^, caused by the hypoxia consecutive to cardiac arrest. As a matter of fact, isoflurane and ketamine, but not pentobarbital, have been proposed as protecting against spreading depolarization^18,19^, which may explain some of the differences observed between groups 1-2 and group 3-4. Moreover, mitochondria respond instantly to sudden changes in cerebral oxygenation and post-mortem hypoxia, even of short duration, leads to fragmentation of mitochondria, as demonstrated in a recent publication^5^. As a matter of fact, these peri-mortem alterations change drastically the baseline of measured parameter and although the implication for data collection might be minimal when comparing a control and experimental condition prepared the same way, these changes are important to consider when switching from one method of tissue fixation to another. This would for example complicate the interpretation of longitudinal studies or comparison with past experiments, and may warrant to redo some previous experiments to generate results in comparable situations. Furthermore, axon fragmentation and smaller size mitochondria in the control condition may make it more difficult to detect subtle phenotypes in the experimental condition, leading to false negative results.

On the contrary, our results show no impact on dendritic spines maturity. One limitation of our study is that it was carried out on in one strain of mouse, looking at one specific brain region and one specific age (P21). We don’t rule out that different results could be observed at different time points where synaptic plasticity is more important, nor that other brain regions would not be affected. However, the quality of the tissues can 1) induce difficulties during the confocal microscopic acquisition and 2) disrupt the analysis if there is so much background noise for example. Note that our study was carried out with wild-type animals, in which dendrites are normally formed. Alterations in the morphology and/or dynamics of dendritic are often associated with neurodegenerative or mental retardation diseases such as FXS^20^. In such pathological models, more variability between animals could have a higher impact and increase bias when interpreting results. It is often assumed that blood removal and perfusion serve the purpose of removing unwanted autofluorescence. Accordingly, we observed more background fluorescence in group 4, although not to such extent as to render imaging uninterpretable. Importantly, although brains from group 3 animals were generally correctly perfused, there were some regions in the brain where small blood vessels remained visible, indicating that fixative agent distribution using an osmotic pump may introduce some variability from one tissue to the other. Although this was generally not consequential, poorly perfused regions sometimes interfered with the analysis, such as for the visualization of CREB activity (**see Fig. 5**). Here, we chose to electroporate neurons with a plasmid coding for the bright red fluorescent protein mScarlet-I, which is one of the brightest fluorescent protein available. mScarlet-I has several advantages: it allows for a good visualization of cellular structures, it remains bright after fixation, and does not require immuno-amplification which could increase the autofluorescence of the tissue. Furthermore, the red channel is usually less impacted by autofluorescence than the green channel. Finally, we imaged dendritic spines with a sensitive Nikon A1 confocal microscope and GaASP detectors. In these optimal imaging conditions, signal to noise ratio is optimal and potential differences in autofluorescence may have a lesser impact. Experimenters would have to adapt to their choice of fluorescent protein and microscope available and verify that in their condition the signal can be accurately distinguished from autofluorescence before adopting a tissue fixation modality.

Handling time varied importantly between groups, with differences in how many times the animal had to be manipulated, which can be source of stress. What’s more, the quality of sedation was variable between the four experimental groups. In our experience, more variability was observed with injectable anesthesia in group 2. Moreover, the delay between the beginning of the procedure and the arrival of the fixing agent in the brain could be very important (especially in group 3). This had a direct impact on the duration of the perfusion per animal and therefore on its well-being, but it also affects the experimenter. As a typical experiment usually involves the preparation of a whole litter of animals, it is not uncommon to prepare 10 to 12 animals at the same time. The increased manipulation and overall preparation time impose a stress on the experimenter that should not be disregarded.

The justification of the use of live animal in experimental neurosciences is that knowledge advance (ie. the benefit of the experiment) outweighs its ethical cost. Aside from the 3R principle from Russel and Burch, it is equally important to generate robust data. Complementary ‘R’ words, including ‘Reproducibility’ or ‘Robustness’ have been proposed in a broader ‘6R’ principle for animal experimentation^21^. Our study falls in line with this movement, by guiding the community toward choosing the best fixation condition. Although we don’t claim that one method is in essence better than the others, we demonstrate that the choice of a fixation method must take into account the expected parameter of analysis, as well as the well-being of the experimenter, and minimizing the suffering of the animal. Thus, our results strongly argue against a one-size-fit-all method that could be uniformly applied to all experiments. Future studies may address other important aspects of this procedure, for example by defining early (and easy to implement) criteria for detecting death, which will make it possible to consider earlier post-mortem perfusions.

## MATERIALS AND METHODS

### Animals

Experiments were carried in respect to the French legislation regarding animal experimentation. Experimenters have FELASA level c (‘conceptor’) certification and justified of training in laboratory animal science in accordance with the EU Directive 2010/63. They also passed a certifying training in surgery. Experimental protocol was approved by the ACCeS Ethics committee. Two pregnant swiss females were purchased from Janvier Labs. They were transported to the animal facility at E7.5 (embryonic day) and were allowed ad libitum access to food, water and maintained on a 12-hr light-dark cycle. In utero cortical electroporation was performed at E15.5.

### DNA and Plasmids

Endotoxin-free plasmid DNA was obtained using the Macherey Nagel midi-prep kit, according to the manufacturer’s instructions. The pCAG-mScarlet-I was described previously^22^. The pCAG-OMM-GFP was kindly provided by Tommy L. Lewis (Oklahoma Medical Research Foundation, USA).

### In utero cortical plasmid electroporation

In Utero cortical plasmid electroporation were performed at E15.5 as described in (Meyer-Dilhet and Courchet, 2020)^6^. A mix containing 1 μg/μl of pCAG-mScarlet-I (to see the neurons), 0.25 μg/μl pCAG-OMM-GFP (to see the mitochondria) and 0.5% Fast Green (Sigma; 1/20 ratio) was injected into one lateral hemisphere of embryos using microcapillaries. Electroporation was performed using an ECM 830 electroporator (BTX) with 3×5mm GenePad electrodes using 4 pulses of 45V during 50msec, with 500 msec interval to target cortical progenitors. Analgesia was achieved by pre-operative injection of Buprenorphine (0.1mg/kg of body weight). Post-surgical pain was monitored for two days using a standardized scale approved by the ethical committee, and analgesia was provided accordingly using Buprenorphine (0.1mg/kg of body weight) for 2 days. Delivery typically occurred at gestational age E19. At postnatal day 2 (P2), we verified the efficiency of electroporation with a Xite Fluorescence flashlight system (Nightsea, distributed by Electron Microscopy Sciences). Non electroporated pups were culled, and 16 pups were conserved. At P21, the 16 pups were allocated to 4 groups in order to mix litter and male/female ratio in each group and were sacrificed using the appropriate procedure.

### Tissue fixation method

All mice received premedication/analgesia: Buprenorphine (Buprenex®) diluted at 0.03 mg/mL (final dose 0.1mg/kg) by subcutaneous injection at least 30 minutes-2h before sacrifice.

In this study, we compared four tissue fixation methods:

- Group 1 and 2: Transcardiac perfusion ante-mortem The mice were anesthetized by volatile anesthesia (Isoflurane 5%, O2 1,5L/min) (group 1) or by fixed anesthesia by an intraperitoneal injection of ketamine (150mg/kg)/xylazine (15mg/kg) (group 2). Once deep sedation was achieved (assessed by the loss of reflex to toe pinching and slowering of the respiration rate), mice were placed on dorsal decubitus. For group 1, mice were maintained under anesthesia under a mask (Isoflurane 5%, O2 1,5L/min). A large incision of the skin revealed the ribcage, which was incised using scissors to expose the heart. The right atrium was incised, then mice were sacrificed by intracardiac perfusion of 1X PBS (total volume 5mL), and then injection of 4% paraformaldehyde/1X PBS (total volume 10mL) (PFA).

- Group 3: Transcardiac perfusion post-mortem

The mice were anesthetized by intraperitoneal injection of xylazine (15mg/kg) to induce sedation. Five minutes after, we performed an intravenous injection of heparin (500UI/kg, 10μL) in the caudal vein. Then, mice were sacrificed by intraperitoneal injection of a veterinary approved euthanizing agent (pentobarbital, 140mg/kg). Death was characterized by the absence of respiratory movement and absence of reflex to toe pinching. As quickly as possible once death has been noted (5-10 min post-injection), mice were placed on dorsal decubitus. A large incision of the skin revealed the ribcage, which was incised using scissors to expose the heart. The right atrium was incised, then 1X PBS was injected in the heart using a peristaltic pump (speed 10mL/min) for 1 minute. Immediately after, a solution of 4% paraformaldehyde/1X PBS (total volume 10mL) was injected using the peristaltic pump (speed 10mL/min) for about 5min.

- Group 4: No perfusion

This group is a negative control to verify that the fixative injections of groups 1 to 3 provide better quality tissue.

The mice were sacrificed by an intraperitoneal injection of xylazine (15mg/kg) to induce sedation, and five minutes later an intraperitoneal injection of veterinary euthanizing agent (pentobarbital, 140mg/kg). Death was characterized by the absence of respiratory movement and absence of reflex to toe pinching. As quickly as possible once death has been noted (5-10 min), the brains of the mice were extracted and dipped into a solution of fixative agent (4% paraformaldehyde/1X PBS).

For all groups, a second series of animals was performed following the exact same procedure but replacing the 4% paraformaldehyde/1X PBS (PFA) solution by a 2% paraformaldehyde/0.075% glutaraldehyde/1X PBS solution (PFA-GA).

All whole brains are post-fixed overnight in PFA or in PFA-GA at 4°C, protected from light.

### Brain slicing and Immunostaining

Following fixation and 1X PBS washing (3 times for 10 min), brains were embedded in 3% low melt agarose in 1X PBS. 80μm thick sections were performed using a Leica VT1000S vibratome (2 slices focus on the corpus callosum and 2 on the hippocampus per animal). Slices were then incubated overnight at 4°C under agitation with primary antibody polyclonal chicken anti-GFP (Rockland) diluted at 1/2000 in permeabilization buffer (PB: 5% BSA, 0.1% TritonX-100 and 1X PBS) protecting from light. The next day, slices were washed 3 times for 10 min in 1X PBS under agitation and then incubated with secondary antibodies (polyclonal goat anti-chicken, Alexa 488, 1/2000 (Life technology). Nuclear DNA was stained using Hoechst 33258 1/5000 (Pierce) 1h at room temperature. After 3 washes in 1X PBS, 10 min at room temperature under agitation, slices were mounted on slides with fluoromount^G^ (Invitrogen) and kept à 4°C.

### Image acquisition and Analyses

#### Axonal projections

Confocal images were acquired in 1024×1024 mode with a Nikon Ti-I microscope equipped with a C2 or a A1 laser scanning confocal microscope.

Microscope control and image analysis was performed using the Nikon software NIS-Elements (Nikon). We used the following objective lenses (Nikon): 10X PlanApo; NA 0.45, 20X PlanApo VC; NA 0.75. The parameters (laser power and gain of the detector) were set for the first series of images in order to avoid saturated signal and conserved, within one series, in order to get signal intensity comparable from one slice to the other.

On the ipsilateral, five ROIs were placed in the corpus callosum (for normalization), and in the cortex (background fluorescence). Signal was defined as (average (layer V) – average (background)) / (average (corpus callosum) – average (background)). To assess terminal axon branching in vivo, we measured mScarlet-I fluorescence intensity of axonal projection on the contralateral side along the six layers of the cortex and normalized to mScarlet-I intensity in the axons of the white matter, which is correlated to electroporation efficiency.

#### Mitochondria and dendritic spines

Images were acquired with Nikon Ti-I microscope equipped with a C2 or a A1 laser scanning confocal microscope. High resolution images of dendritic spines and mitochondria from pCAG-mScarlet-I / pCAG-GFP-OMM mix electroporated neurons were acquired using the 100X (N.A.: 1.45; zoom:1.5; resolution: 2048×2048 px). Z stacks (z-step:0,15μm) of digital images defined from top to bottom were captured using the Nikon software NIS-Elements (Nikon). For the figures, the Z stacks were collapsed in one resulting picture using the maximum intensity projection function provided by Fiji (Image J) software. Analysis of mitochondrial length and occupancy as well as dendritic spines density and morphology were performed using Fiji.

### Quantifications and Statistical analyses

Statistical analysis were performed using GraphPad Prism. The parameters measured (axonal fragmentation, atrophy of synapses, fragmentation of mitochondria) were compared by a Student’s t test to compare two experimental groups or an ANOVA test to compare 3 experimental groups or more.

Quantifications were performed blind to experimental condition.

Statistical significance is indicated in all figures by the following annotations: ns, not significant ; *, p<0.05; **, p<0.01 ; ***, p<0.001.

## ACKNOWLEDGEMENTS

The authors thank members of the Courchet lab and Institut NeuroMyoGène for useful comments and discussion. We thank Laura Barrot and Stephan Langonnet for suggestions and comments. We are grateful to members of the Ethical committee ACCeS for critical assessment of the study design. We thank the staff of the SCAR and P-PAC facilities for assistance in conducting this study. S.E. was recipient of a postdoctoral fellowship from LabEx Cortex.

## AUTHORS CONTRIBUTION

G.M.D. and J.C. conceived the study, performed in utero cortical electroporations and tissue preparation. G.M.D. prepared tissues and histological samples, quantified and interpreted experiments for axonal preparations and immunohistochemistry. S.E. quantified and interpreted experiments for mitochondria morphology and dendritic spines density and morphology. O.R. provided critical insight on study design, and analyzed immunohistochemistry with G.M.D. J.C. supervised the work. J.C. and G.M.D. drafted the manuscript and all other authors edited and approved the manuscript.

