## Supplementary figures for "Comparative study of pre- and post-mortem perfusion of fixative for the quality of neuronal tissue preparation"

### TITLE

#### SUPPLEMENTARY FIGURES

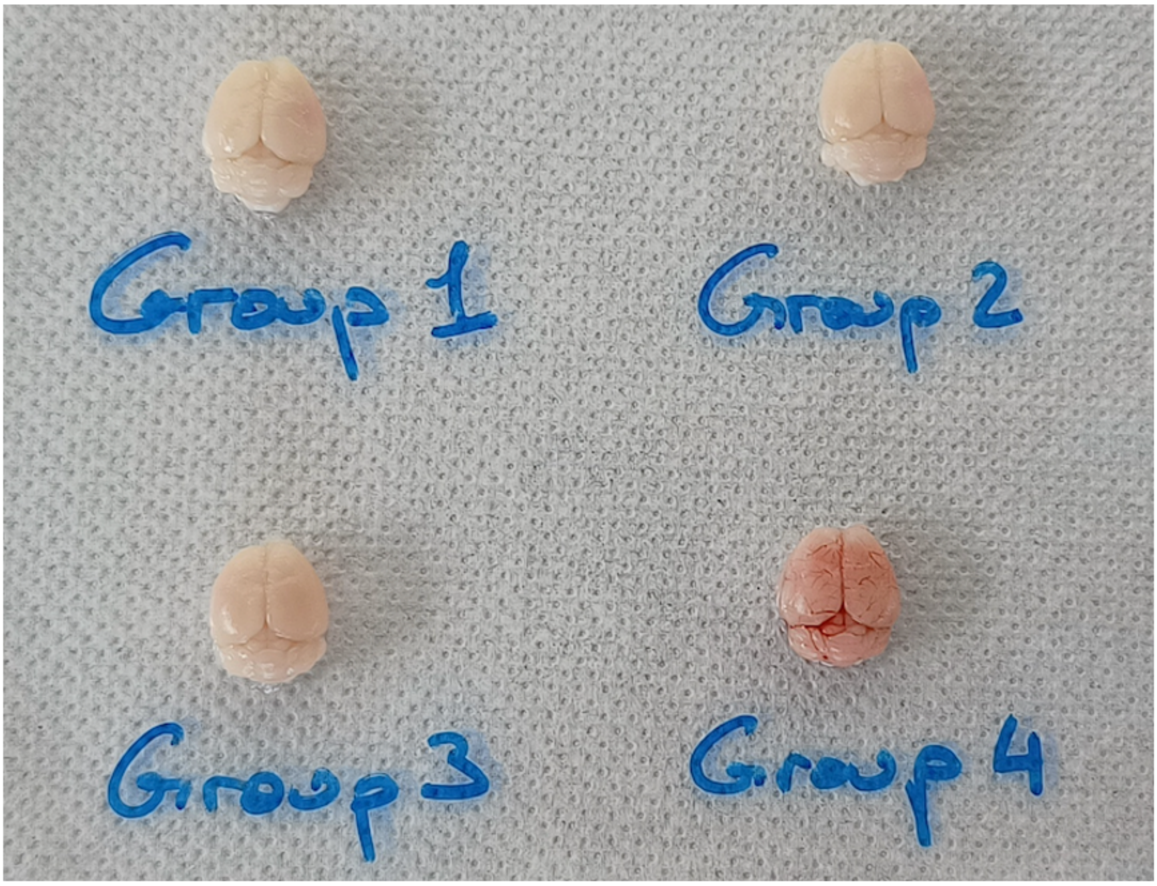


**Supplementary Figure 1: Macroscopic aspect of the brains following dissection**

Photograph of brains representative of the perfusion quality for each group. Brains were washed in PBS for the photography, but not yet dipped into fixative.

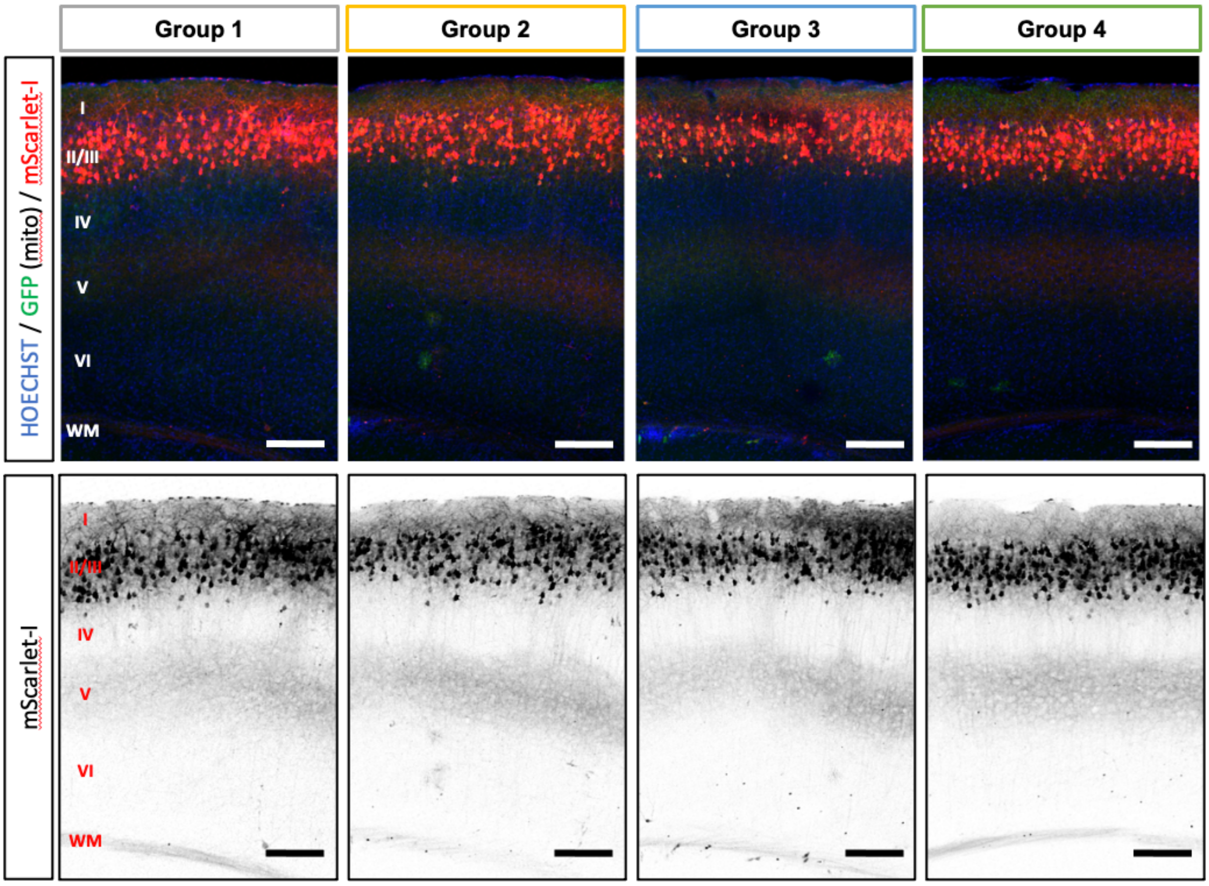


**Supplementary figure 2: Detail of the ipsilateral cortex for each group**

Representative images of the ipsilateral side for each group. Top: overlay of HOECHST, GFP (marking mitochondria) and mScarlet-I (marking the soma and axons). Bottom: mScarlet-I signal. Scale bar: 200µm. WM: white matter. I, II, III, IV, V and VI: cortical layer identity.

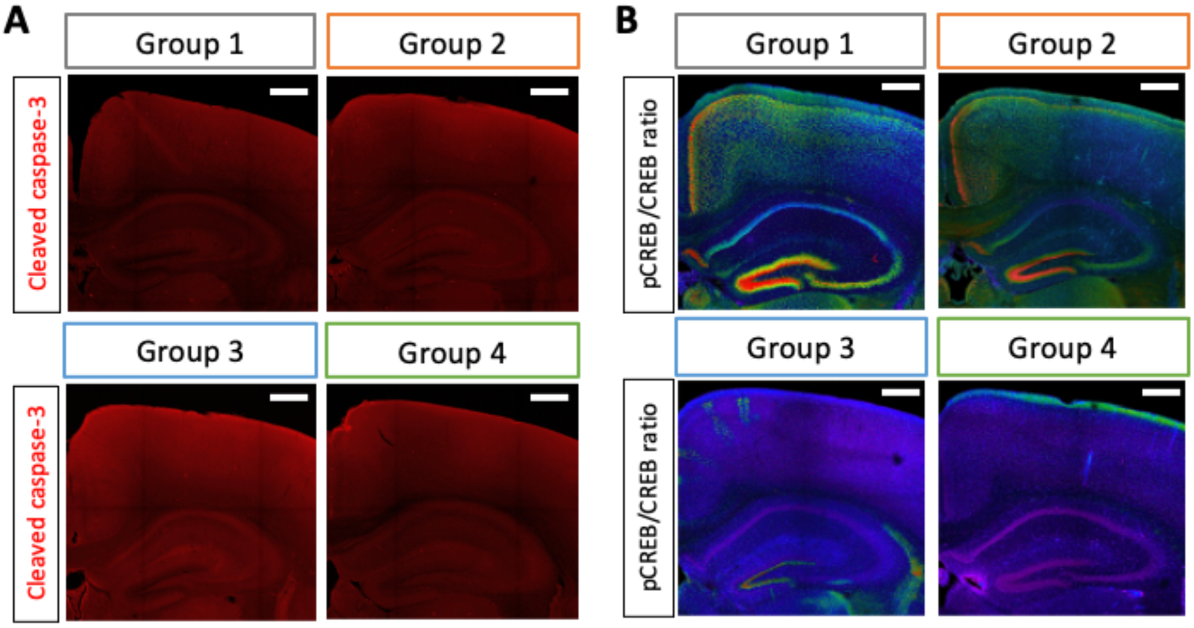


**Supplementary Figure 3: Effect of perfusion condition on immunostaining upon PFA-GA fixation**

**(A)** Neuronal apoptosis observed by cleaved caspase-3 staining in each group, following perfusion/fixation with PFA. **(B)** Ratio between phosphorylated CREB (pCREB, Ser133) and non phosphorylated CREB in each group. Images are colorized on a rainbow LUT (purple: low ratio. red: high ratio). Magnification of the cortex and hippocampus. Raw images for CREB and pCREB provided in **Suppl. Fig. 4**. Scale bar: 500µm.

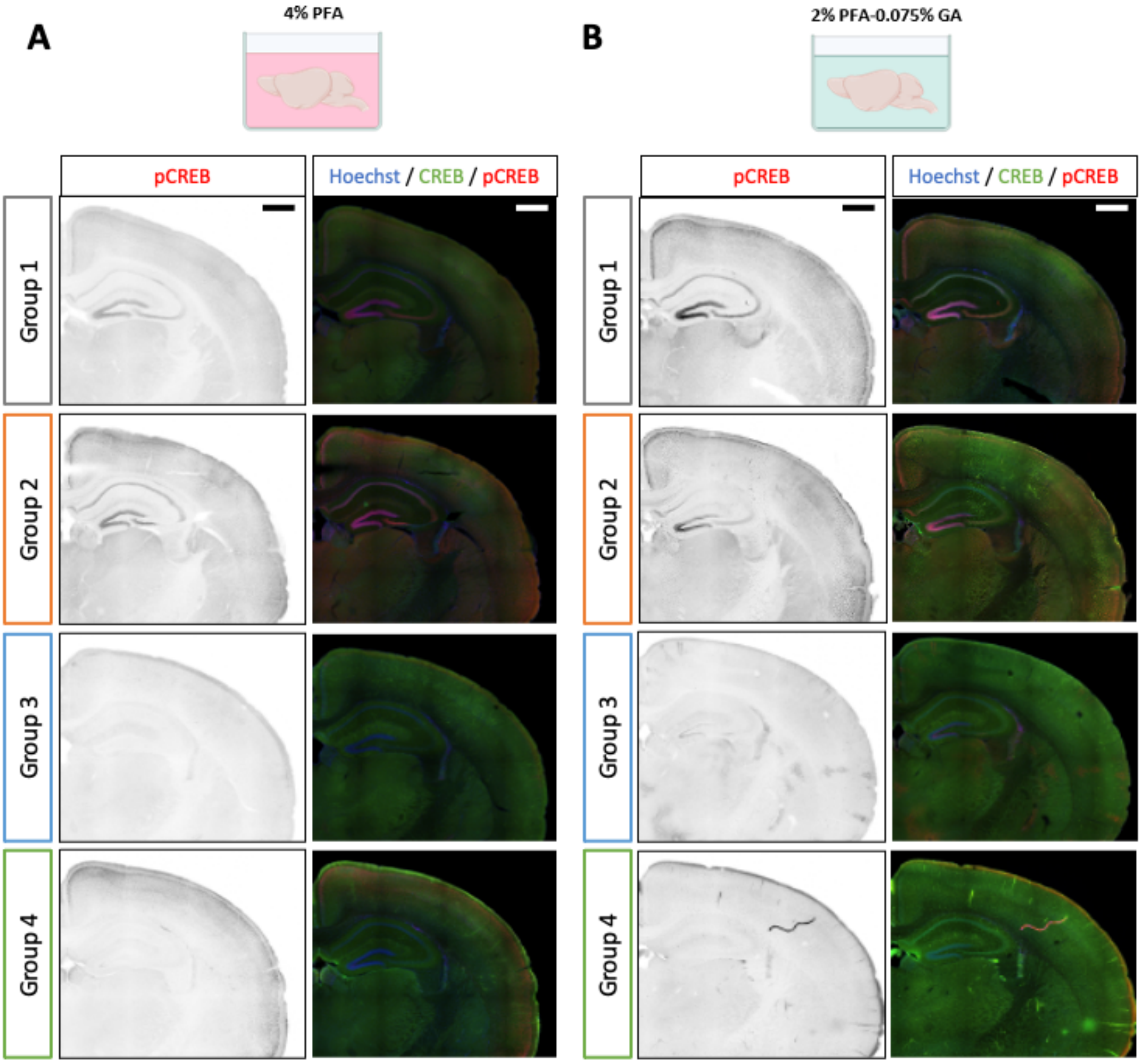


**Supplementary figure 4: Raw images of pCREB and CREB immunostaining**

Representative coronal sections of the immunoreactivity of pCREB (Serine 133) and CREB in each group, for fixation with PFA **(A)** or PFA-GA **(B)**. Hoechst was used to stain nuclei and provide brain architecture. pCREB staining is presented as a mono-channel image and inverted color. Scale bar 1mm
